# Transdiagnostic Neurobiology of Social Cognition and Individual Variability as Measured by Fractional Amplitude of Low-Frequency Fluctuation in Schizophrenia and Autism Spectrum Disorders

**DOI:** 10.1101/2024.07.02.601737

**Authors:** Soroush Bagheri, Ju-Chi Yu, Julia Gallucci, Vinh Tan, Lindsay D. Oliver, Erin W. Dickie, Ayesha G. Rashidi, George Foussias, Meng-Chuan Lai, Robert W. Buchanan, Anil K. Malhotra, Aristotle N. Voineskos, Stephanie H. Ameis, Colin Hawco

**Author notes:** **Corresponding Author:** Dr. Colin Hawco, PhD, Assistant Professor, Department of Psychiatry, University of Toronto. Scientist, Brain Health Imaging Centre, Centre for Addiction and Mental Health. Room 623, 250 College Street, Toronto, ON. M5T 1R8, (P). Equal contribution.

## Abstract

Fractional amplitude of low-frequency fluctuation (fALFF) is a validated measure of resting-state spontaneous brain activity. Previous fALFF findings in autism and schizophrenia spectrum disorders (ASDs and SSDs) have been highly heterogeneous. We aimed to use fALFF in a large sample of typically developing control (TDC), ASD and SSD participants to explore group differences and relationships with inter-individual variability of fALFF maps and social cognition. fALFF from 495 participants (185 TDC, 68 ASD, and 242 SSD) was computed using functional magnetic resonance imaging as signal power within two frequency bands (i.e., slow-4 and slow-5), normalized by the power in the remaining frequency spectrum. Permutation analysis of linear models was employed to investigate the relationship of fALFF with diagnostic groups, higher-level social cognition, and lower-level social cognition. Each participant’s average distance of fALFF map to all others was defined as a variability score, with higher scores indicating less typical maps. Lower fALFF in the visual and higher fALFF in the frontal regions were found in both SSD and ASD participants compared with TDCs. Limited differences were observed between ASD and SSD participants in the cuneus regions only. Associations between slow-4 fALFF and higher-level social cognitive scores across the whole sample were observed in the lateral occipitotemporal and temporoparietal junction. Individual variability within the ASD and SSD groups was also significantly higher compared with TDC. Similar patterns of fALFF and individual variability in ASD and SSD suggest some common neurobiological deficits across these related heterogeneous conditions.

## 1. Introduction

Autism spectrum disorders (ASD) and schizophrenia spectrum disorders (SSD) are phenomenologically similar disorders, which affect approximately 1% of the population [1,2]. Exploring spontaneous brain activity of people with these diagnoses in a resting-state condition, where brain activity is not affected by the effects of various tasks, could provide more insight into the inherent neurobiological deficits of these disorders. The amplitude of low-frequency fluctuations (ALFF) and fractional ALFF (fALFF) are derived from the examination of functional magnetic resonance imaging (fMRI) blood oxygen level-dependent (BOLD) signal power in the low-frequency ranges (0.01 to 0.1 Hz); they have the benefit of allowing for examining task-free, resting-state spontaneous local brain activity in healthy and clinical populations. In addition, fALFF, compared with ALFF, has the advantage of accounting for power across the full frequency range and may be less susceptible to physiological noise [3,4].

Investigation of ALFF and fALFF in SSD and ASD, have yielded heterogeneous results. For instance, in some studies, SSD has shown increased resting-state activity in medial prefrontal cortex (mPFC) [5,6], precuneus [7] and insula [8,9], while other studies have shown decreased (f)ALFF in the same regions: mPFC [10], precuneus [9–11], and insula [5]. A meta-analysis comparing ALFF in SSD with typically developing controls (TDC) found increased ALFF in the right putamen, left inferior temporal gyrus, and right inferior frontal gyrus, along with decreased ALFF in postcentral gyrus and left precuneus. However, this meta-analysis also noted substantial heterogeneity among findings, which was attributed to the methodological diversity across studies [12]. Fewer studies have examined (f)ALFF in ASD; one study found higher activity in ASD than TDC in the right middle temporal gyrus, inferior parietal gyrus, and angular gyrus [13]. However, lower (f)ALFF in ASD versus TDC has also recently been observed in the left parietal opercular cortex, right insular cortex, and precuneus [14].

An important feature of both ASD and SSD is atypical and/or impaired social cognition, which can be associated with poor functional outcomes [15]. Social cognition refers to the capability to perceive, process, and respond to social stimuli [16]. Recent work has suggested that, generally, social cognitive processes in SSD can be divided into lower-level processes, such as emotion recognition, and higher-level processes, such as detecting sarcasm [17,18]. A meta-analysis has revealed that SSD and ASD show similar levels of impairment relative to TDC in both lower- and higher-level social cognitive processes [19]. A few studies of (f)ALFF have found associations with social cognitive deficits in either schizophrenia or autism. For example, a multi-modal biomarker study of cognition in SSD noted that higher fALFF in the inferior parietal lobule and angular gyrus was strongly correlated with poorer social cognition [20]. Another study in early-onset schizophrenia found a negative correlation between theory of mind sense of reality and ALFF in the left precuneus [21]. Also, in boys with autism, fALFF in the right lateral occipital cortex was negatively correlated with social communication scores [22].

Recent work from our group has highlighted increased inter-individual variability in SSD and ASD in task-induced fMRI activity [23,24]. Inter-individual variability is a metric that quantifies how “typical”, or close to the average, one’s pattern of brain activity is. A study that examined a spatial working memory N-Back task showed greater individual variability in the spatial pattern of brain activity in ASD [23], demonstrating that individuals with ASD were more likely to show “idiosyncratic/atypical” brain activity during cognitive processing. This increased variability was also shown in SSD during a letter-sequence N-Back working memory task [24], a finding that was subsequently demonstrated to be greater in SSD than bipolar disorder, and related to illness duration in SSD only [25]. This increased individual variability may represent an important diagnostic marker that is not well-captured by group comparisons [23,25].

In the current study, we aimed to extend the literature in three ways: (1) by examining fALFF across a large transdiagnostic sample including individuals with SSD or ASD, and TDCs, allowing for groupwise comparisons between ASD and SSD, (2) by relating fALFF to both lower- and higher-level social cognition tasks, and (3) by examining fALFF-derived individual variability, in an effort to understand inter-individual patterns in fALFF-measured local brain activity. Data was examined from the Social Processes Initiative in the Neurobiology of Schizophrenia(s) (SPINS) and the Social Processes Initiative in the Neurobiology of Autism Spectrum Disorder (SPIN-ASD), which include social, neurocognitive and resting-state fMRI data [26,27]. Due to the heterogeneous findings of prior studies [12], we hypothesized that the ASD and SSD groups would show differences in fALFF compared with TDCs with no prediction about the region and directionality of these differences.

Furthermore, we hypothesized that fALFF would relate to higher- and lower-level social cognitive scores similarly across groups with common neural circuitry [27], and both SSD and ASD would have higher variability than TDCs [23,25].

## 2. Methods and Materials

### 2.1 Participants

Eligible participants (total *N* = 578; 75 ASD, 301 SSD, 202 TDC) were recruited from three different institutions: the Centre for Addiction and Mental Health (CAMH; Toronto, Canada), Maryland Psychiatric Research Centre (MPRC; Maryland, USA), and Zucker Hillside Hospital (ZHH; New York, USA), for the SPINS dataset and one institution (CAMH) for the SPIN-ASD dataset. SPINS and SPIN-ASD are two harmonized, multimodal, National Institute of Mental Health-funded datasets designed to assess brain-behavior associations related to social cognition across individuals with ASD, SSD or TDCs. This study was approved by the institutional review board of each institution and all participants provided informed consent prior to participation. All participants were evaluated using the Structured Clinical Interview for the Diagnostic and Statistical Manual of Mental Disorders (SCID-IV-TR).

Inclusion criteria for participants with ASD included: having a clinical diagnosis of autism based on the Diagnostic and Statistical Manual of Mental Disorders, 5th edition (DSM-5), further confirmed using the Autism Diagnostic Observation Schedule-2 (ADOS-2) [28], having an IQ>70 (i.e., the absence of co-occurring intellectual disability) estimated using the Wechsler Abbreviated Scale of Intelligence (WASI-II), and stable antipsychotic medication prescriptions (and doses) for at least 30 days prior to study enrollment. Inclusion criteria for participants with SSD included: confirmed diagnosis using the DSM-5 for schizophrenia, schizoaffective disorder, schizophreniform disorder, delusional disorder, or psychotic disorder not otherwise specified, IQ>70 as estimated by the Wechsler test of Adult Reading (WTAR) [29], stable doses of antipsychotic medication and no decline in functioning and support level in the 30 days prior to enrollment. For the TDC group, the inclusion criteria were the absence of a current or past axis I psychiatric disorder, phobic disorder, and past major depressive disorder over the previous two years with the exception of adjustment disorder. Furthermore, the TDC group from the SPINS dataset did not have a first-degree relative with a history of a primary psychotic disorder and had to be presently unmedicated. Additional exclusion criteria for all participants included any MRI contraindications, history of head trauma resulting in unconsciousness, diagnosis of a substance use disorder (confirmed by meeting the DSM-5 criteria and/or urine toxicology screening), intellectual disability, any debilitating or unstable medical illnesses, other neurological diseases, and pregnancy (or potential pregnancy). All participants also had normal or corrected-to-normal vision.

### 2.2 Clinical and Cognitive Assessments

Psychiatric symptoms were evaluated using the Brief Psychiatric Rating Scale (BPRS) [30]. The SSD group completed a modified version of the Scale for the Assessment of Negative Symptoms (SANS) [31,32]. The lower-level social cognitive test consisted of the Penn Emotion Recognition Task (ER40) [33], which assesses facial emotion recognition measured by the accuracy of the responses. For a measure of higher-level social cognitive processing, we chose The Awareness of Social Inference Test, Revised Part 3 Sarcasm sub-score (TASIT-3-Sar), which involves viewing videos that depict sarcastic social situations. This score was selected because it has been associated with theory of mind [34,35], and loads well on a higher-level mentalizing factor in SSD [17].

### 2.3 MRI Data Acquisition

SSD data was from a previously collected sample, SPINS [26,27,36]. SPINS participants were scanned across three sites, each of which upgraded MRI scanners part way through, meaning six 3T scanners, with harmonized acquisition parameters [26,37]. From the final sample (*N* = 495), CAMH used a General Electric Discovery (*N* = 133) and a Siemens Prisma (*N* = 103), MPRC used a Siemens Tim Trio (*N* = 64) and a Siemens Prisma (*N* = 73), and ZHH used a General Electric Signa (*N* = 41) and Siemens Prisma (*N* = 81). The SPIN-ASD sample was collected on a Siemens Prisma scanner using an imaging protocol harmonized to the SPINS sample. Relevant scans for the current analysis include a T1-weighted (T1w) anatomical scan (voxel-size = 0.9 mm; see site-specific differences in [37]), as well as a seven-minute resting-state scan using an Echo planar imaging sequence (TR = 2000 ms, TE = 30 ms, flip angle = 77°, voxel size 3.125 x 3.125 x 4 mm, 212 volumes).

### 2.4 fMRI Preprocessing

All structural and functional data were preprocessed using fMRIPrep version 1.5.8 and ciftify [38]. The first 3 TRs of all scans were removed, and all scans were slice-time corrected. A transform into MNI space was calculated and the fMRI data were projected to the cortical surface [38]. Following that, the regression model applied to the data included correction for all head motion parameters (i.e., translation and rotation along the x, y, and z axes), a 2nd order detrending, regressing mean white matter (WM), mean cerebral spinal fluid (CSF) and global signals, as well as the square, the derivative, and the squared derivative for each of these regressors. Surface and subcortical data were then smoothed with a 6mm FWHM gaussian kernel.

### 2.5 fALFF Measurement

fALFF was measured by conducting a Fast Fourier Transform on each voxel time series to convert the signal to a frequency domain power spectrum. fALFF was calculated by dividing the mean square root of the BOLD signal power of each voxel in the frequency range 0.01 - 0.027 Hz for slow-5, and 0.027 - 0.073 Hz for slow-4, by the total power in the full frequency range (0.00 - 0.25 Hz). This spectrum-specific analysis was conducted due to the prior findings that different brain structures are sensitive to various frequency ranges [39,40]. fALFF values were normalized for each participant by dividing the fALFF value of each voxel by the mean fALFF of the entire brain for that participant.

### 2.6 Multi-Site Harmonization and Permutation Analysis of Linear Models

ComBat harmonization was performed in MATLAB R2023a to remove the non-biological effects of the six different scanners from the data [41,42], separately for slow-4 and slow-5 bands. For each frequency band, a contrast of fALFF maps was computed to evaluate pairwise differences in fALFF values; ASD versus TDC, SSD versus TDC, and ASD versus SSD. Slow-4 and slow-5 group contrast maps were analyzed with FSL’s Permutation Analysis of Linear Models (PALM), using threshold-free cluster enhancement (TFCE) across 1000 permutations [43]. The results were then thresholded with Family-Wise Error (FWE) at p< 0.025 to correct for both hemispheres. Mean framewise displacement (FD) [44], age, sex, and scanners were included as covariates. TASIT-3-Sar and ER40 were included as regressors to consider social cognition and account for differences in fALFF based on social cognition. In addition, a group by TASIT-3-Sar interaction, and a group by ER40 interaction analysis were also performed. Another model without social cognitive variables was run to examine if diagnostic differences were present when social cognition was not accounted for. A post-hoc sensitivity analysis was also conducted with an age- and sex-matched subset, and only including participants with Siemens Prisma MRI data. This subsample analysis aimed to retain as many ASD participants as possible (as the smallest group), only across Prisma scanners as all ASD participants were collected on a Prisma MRI, and including TASIT-3-Sar and ER40 as covariates.

### 2.7 Individual variability in fALFF maps via correlational distance

Individual variability was calculated for fALFF maps (separately for slow-4 and slow-5). Each participant’s fALFF maps for all cortical vertices were loaded into RStudio v1-554 as a numeric vector; generating a spatial series with each point corresponding to the fALFF value at one vertex for that individual. Pairwise correlational distances were calculated, quantifying the similarity in the spatial pattern of fALFF across each pair of participants. Individual variability was defined as the mean correlational distance (MCD) of a given participant relative to all others, specifying a single value for each participant [23]; lower MCDs are indicative of participants with spatial patterns of fALFF more similar to the overall group, and higher MCDs are indicative of less ‘typical’ and more idiosyncratic spatial patterns of fALFF values. A linear regression assessed the relationship between individual variability and potential explanatory variables or covariates (diagnostic group, age, sex, TASIT-3-Sar and ER40, site, FD and the interaction between social cognitive scores and group).

### 2.8 Code Availability

Code for the study analyses can be found on https://github.com/TIGRLab/falff_paper. The SPINS dataset is available on the NDA at https://nda.nih.gov/edit_collection.html?id=2098, SPIN-ASD at https://nda.nih.gov/edit_collection.html?id=2923.

## 3. Results

### 3.1 Participants Characteristics

Following exclusion, the final sample size was 185 TDC, 68 ASD, and 242 SSD with complete social cognitive and MRI data available (see consort diagram, Supplemental Figure 1). Participant demographic information and clinical and social cognitive characteristics are presented in Table 1.

**Table 1:** Participant demographics and clinical and social cognitive characteristics. Where appropriate, variables are displayed as mean (standard deviation). All statistical comparisons were conducted using ANOVA except for the comparison of sex which was conducted using a chi-squared test. Where participant data was incomplete for a measure, the number of participants for whom the data was available is noted as *n*. TDC = typically developing controls, ASD = autism spectrum disorder, SSD = schizophrenia spectrum disorder, FD = framewise displacement, WTAR = Wechsler Test for Adult Reading, WASI = Wechsler Abbreviated Scale of Intelligence, BSFS = Birchwood Social Functioning Score, BPRS = Brief Psychiatric Rating Scale, SANS = Scale for the Assessment of Negative Symptoms, ADOS = Autism Diagnostic Observation Schedule, CPZ = Chlorpromazine, TASIT-3 = The Awareness of Social Inference Test-Revised, ER40 = Penn Emotion Recognition Task

| Variables | TDC (n = 185) | ASD (n = 68) | SSD (n = 242) | Group Comparison (p-value) |
| --- | --- | --- | --- | --- |
| Age in years | 31.37 (10.19) | 20.96 (3.94) | 31.73 (9.81) | SSD TDC > ASD<br>6.23e-16 |
| Male: Female | 100:85 | 43:25 | 167:75 | 0.0066 |
| Education in years | 15.83 (2.04) | 12.62 (2.07) | 13.59 (2.15) | TDC > SSD > ASD<br><2e-16 |
| Motion (FD) | 0.1372 (0.0688) | 0.1317 (0.0619) | 0.1495 (0.0887) | TDC ASD SSD<br>0.13 |
| WTAR Standard Score † | 113.50 (10.96)<br>(n = 115) | - | 107.46 (13.70)<br>(n = 162) | TDC > SSD<br>0.00011 |
| WASI Score † | 113.69 (9.169)<br>(n = 13) | 117.59 (12.88)<br>(n = 59) | - | TDC ASD<br>0.305 |
| BSFS Score | 175.21 (18.94) | 126.93 (25.64) | 135.91 (23.82) | TDC > SSD > ASD<br><2e-16 |
| BPRS Score Total | - | 27.46 (5.04) | 31.36 (7.95) | SSD > ASD<br>0.00015 |
| BPRS Positive Symptoms | - | 4.63 (1.16) | 7.89 (3.86) | SSD > ASD<br>3.62e-11 |
| BPRS Anxiety and Depression | - | 8.47 (3.42) | 7.67 (3.46) | ASD SSD<br>0.094 |
| SANS Total Score | - | - | 25.19 (12.31) | - |
| ADOS-2 Clinical Severity Score | - | 6.31 (2.25) | - | - |
| CPZ Equivalents (mg/day)<br>[45] | - | - | 476.22 (395.30)<br>(n = 224) | - |
| TASIT-3-Sarcasm Score | 27.6 (3.44) | 26.99 (3.58) | 23.31 (5.26) | TDC ASD > SSD<br><2e-16 |
| ER40 Score | 33.62 (3.22) | 32.29 (4.13) | 31.78 (4.51) | ASD TDC > SSD<br>1.86e-5 |

† SPINS used the WTAR and SPIN-ASD used the WASI.

### 3.2 Group differences in fALFF across TDC, ASD, and SSD

In the model that included TASIT-3-Sar and ER40, reduced fALFF in the visual regions (specifically occipital cortex and cuneus) were observed in the ASD and SSD groups in both slow-4 and slow-5 bands compared with TDC (Figure 1). Compared with TDC, the SSD group also had reduced fALFF in motor regions, and increased fALFF in subcortical regions (hippocampus, amygdala, caudate, and striatum). Significantly higher slow-4, but not slow-5, fALFF were observed in the cuneus regions of individuals with SSD than ASD. Increased slow-5 fALFF was observed in the frontal regions for both ASD and SSD versus TDC, with the ASD group showing more widespread effects. Similar patterns were found when neither TASIT-3-Sar or ER40 was included as covariates in the model (Supplemental Figure 2).

**Figure 1:**
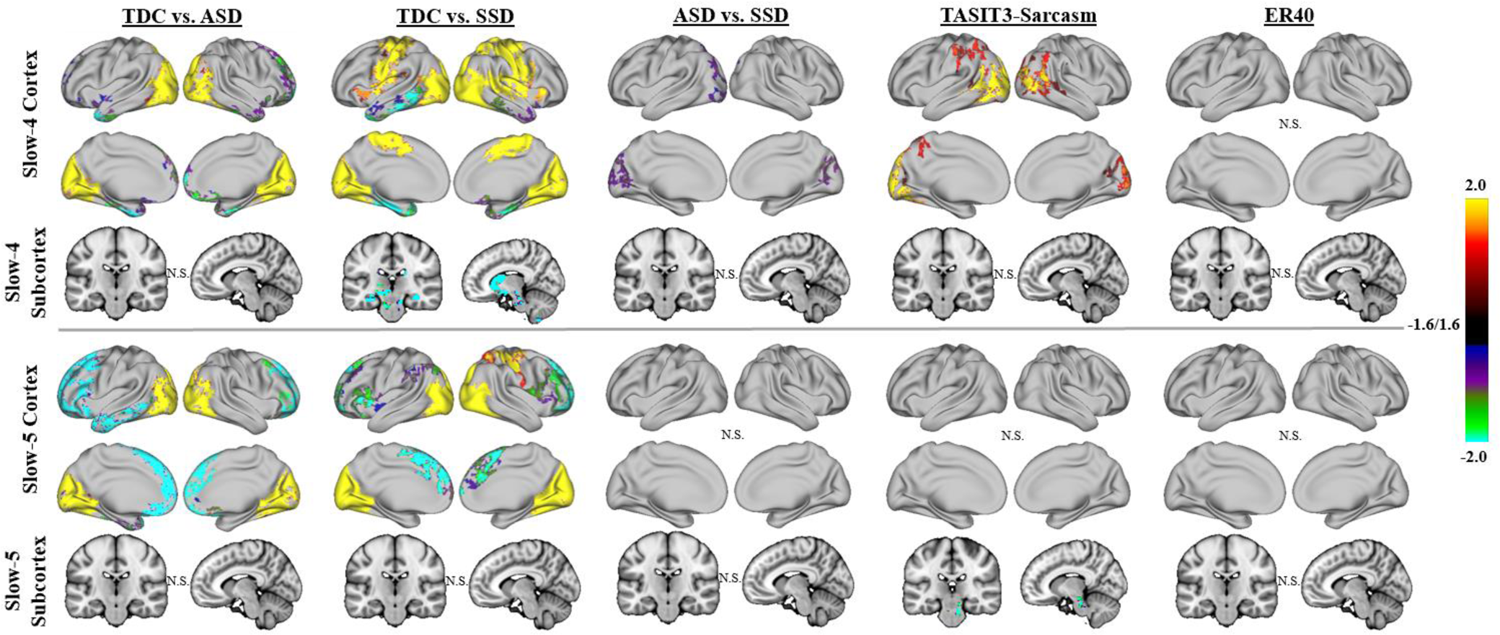
Group results for linear models examining fALFF across the full sample of ASD, SSD, and TDC. Results of whole-brain, vertex-wise analysis using threshold-free cluster enhancement (TFCE) are presented. Results are shown in brain-wide FWE corrected log-10(p), thresholded at 1.6, equal to p = 0.025 (to correct for left and right hemispheres). Significant differences were noted with TDC showing higher fALFF than ASD and SSD in both slow-4 and slow-5 in the visual regions (i.e. cuneus and occipital cortex; yellow-red regions); SSD also had lower fALFF than TDC in the sensory-motor cortex (yellow-red regions). Increased fALFF was observed in the SSD and ASD compared with TDC in the medial frontal regions (green-blue). SSD showed greater fALFF than TDC in the subcortical regions for slow-4 (green-blue). Higher fALFF in SSD compared with ASD was observed in the slow-4 band in the cuneus regions (purple). TASIT-3-Sar scores across the whole sample were positively associated with fALFF in the lateral occipitotemporal and temporoparietal junction in slow-4 (yellow-red) and negatively associated with slow-5 fALFF in the brainstem (green-blue). No significant association of fALFF was found with ER40 scores.

### 3.3 Relationship of fALFF with TASIT-3-Sar and ER40

Across groups, better TASIT-3-Sar scores were associated with higher slow-4 fALFF in the lateral occipitotemporal and temporoparietal junction, and with lower slow-5 fALFF in the brainstem (Figure 1). No statistically significant associations were observed between fALFF across groups and ER40 scores in either cortical or subcortical regions. A significant group by ER40 interaction in both the slow-4 and slow-5 bands was observed for ASD versus SSD and TDC versus ASD in the cuneus regions. In these regions, as ER40 scores increased, fALFF values decreased in ASD but increased in SSD and TDC (Figure 2). Our findings did not indicate a group by TASIT-3-Sar interaction.

**Figure 2:**
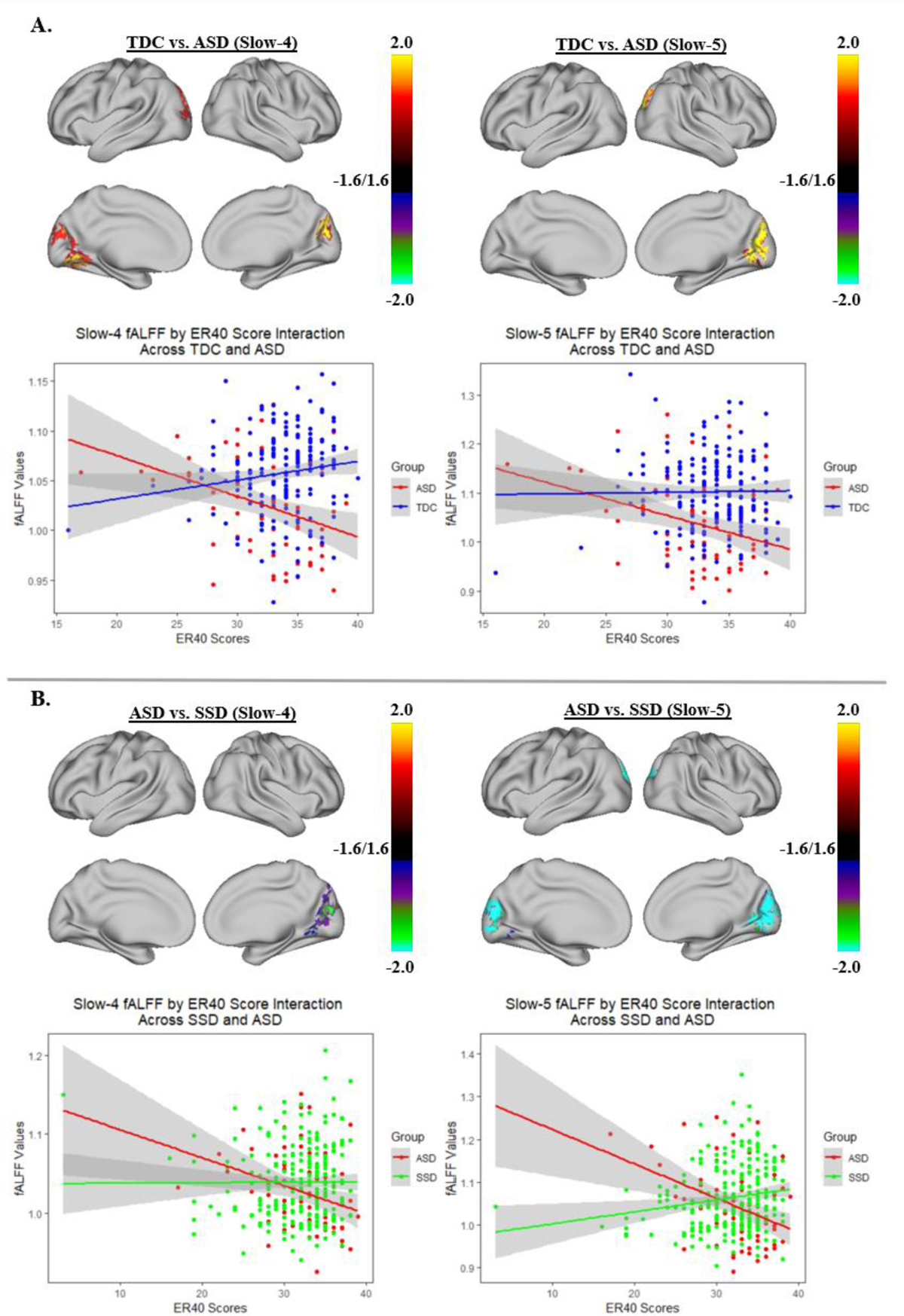
Group by ER40 score interaction effect for ASD compared with TDC (A) and ASD compared with SSD (B). Each of the two panels shows the areas of the cortex where there was a group by ER40 score interaction for slow 4 or slow 5 as well as a scatter plot of the relationship of fALFF and ER40 scores in the highlighted cortical regions. Overall, as ER40 scores increased, both slow 4 and slow 5 fALFF values tend to decrease in the cuneus regions for ASD but increase for SSD and TDC.

### 3.4 Age and Sex Matched Subsample Post-hoc Analysis

After matching the three groups for age and sex (ASD: *n* = 41, mean age = 22.95, 15 female; SSD: *n* = 40, mean age = 24.05, 13 female; TDC: *n* = 43, mean age = 24.26, 21 female; one-way ANOVA for age, p = 0.186, F = 1.705, degrees of freedom (df) = 2; Pearson’s chi-squared test for sex, p = 0.282, *X*^2^, df = 2), the overall pattern of differences in fALFF between ASD or SSD and TDC were similar, but more spatially constrained (Figure 3). ASD showed reduced fALFF in the visual regions in both slow-4 and slow-5. SSD showed reductions in motor regions and insula in slow-4, and visual regions in slow-5. Moreover, SSD had higher fALFF than ASD in the bilateral cuneus and lateral occipital regions in slow-4. No significant differences were observed between ASD and SSD in slow-5. The matched subsample also showed a more spatially constrained association of slow-4 fALFF across groups with TASIT-3-Sar. Moreover, age and sex matching led to some association of slow-4 fALFF with ER40 in the cuneus regions (Figure 3).

**Figure 3:**
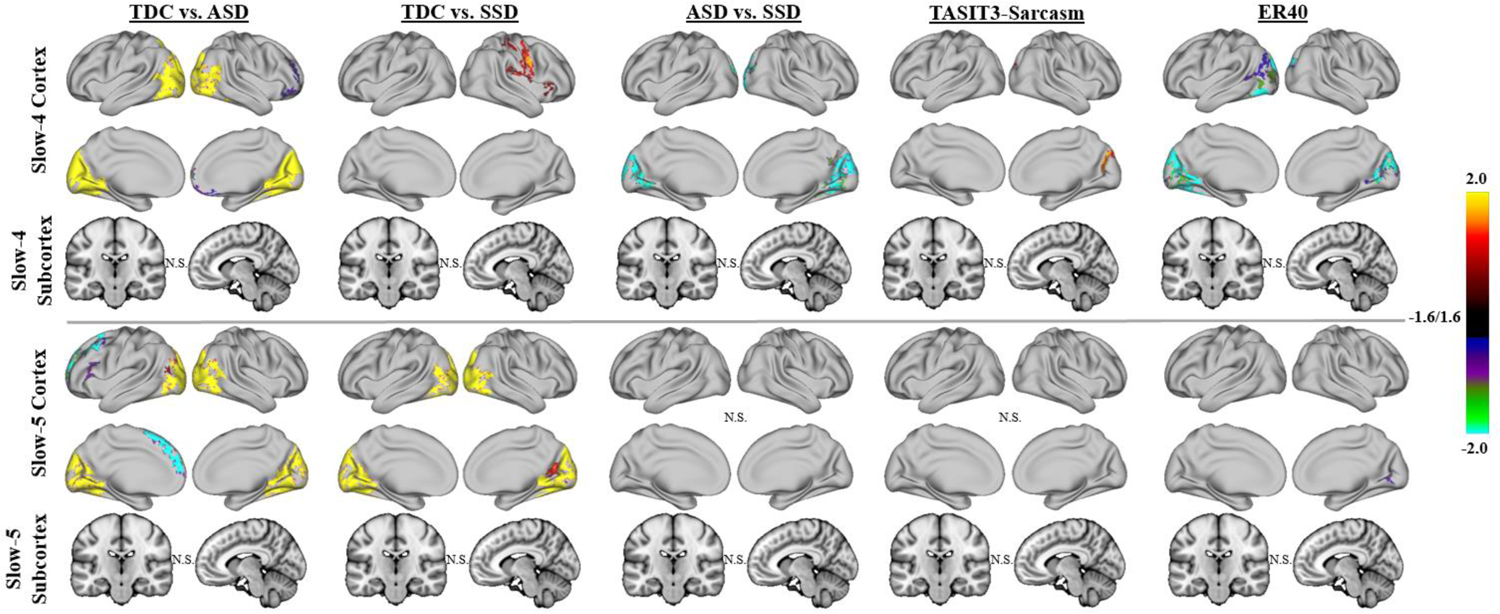
Group results for linear models examining fALFF across ASD, SSD, and TDC, using an age and sex-matched subsample. Higher fALFF in the visual regions in TDC compared with ASD largely persisted after matching the data for age and sex. These differences compared with SSD also persisted in slow-5 but not slow-4. In addition, after matching the data for age and sex, differences between SSD and ASD persisted whereas the association of TASIT-3-Sar and fALFF became more spatially limited. Some association of slow-4 fALFF with ER40 was also observed in the cuneus regions after matching the participants by age and sex.

### 3.5 Mean Correlational Distance Analysis

#### 3.5.1 fALFF slow-4

After accounting for mean FD (F(1,481) = 9.24, p = 0.003) and site effects (F(2,481) = 1.18, p = 0.31), MCD of fALFF slow-4 maps across SSD (0.61 ± 0.02) and ASD (0.61 ± 0.02) were significantly higher than TDC (0.60 ± 0.02), F(2,481) = 10.03, p < 0.001; TDC demonstrated more ‘typical’ spatial patterns of fALFF slow-4, whereas SSD and ASD showed a greater deviation from the norm on average (Figure 4A). Additionally, older age was related to increased MCD, F(1,481) = 26.95, p < 0.001 (Figure 4B), and lower TASIT-3-Sar score was associated with increased MCD, F(1,481) = 12.94, p < 0.001 (Figure 4C). Notably, the effect of social cognition was not influenced by the diagnostic group (diagnosis by TASIT-3-Sar interaction, F(2,481) = 2.06, p = 0.13).

**Figure 4:**
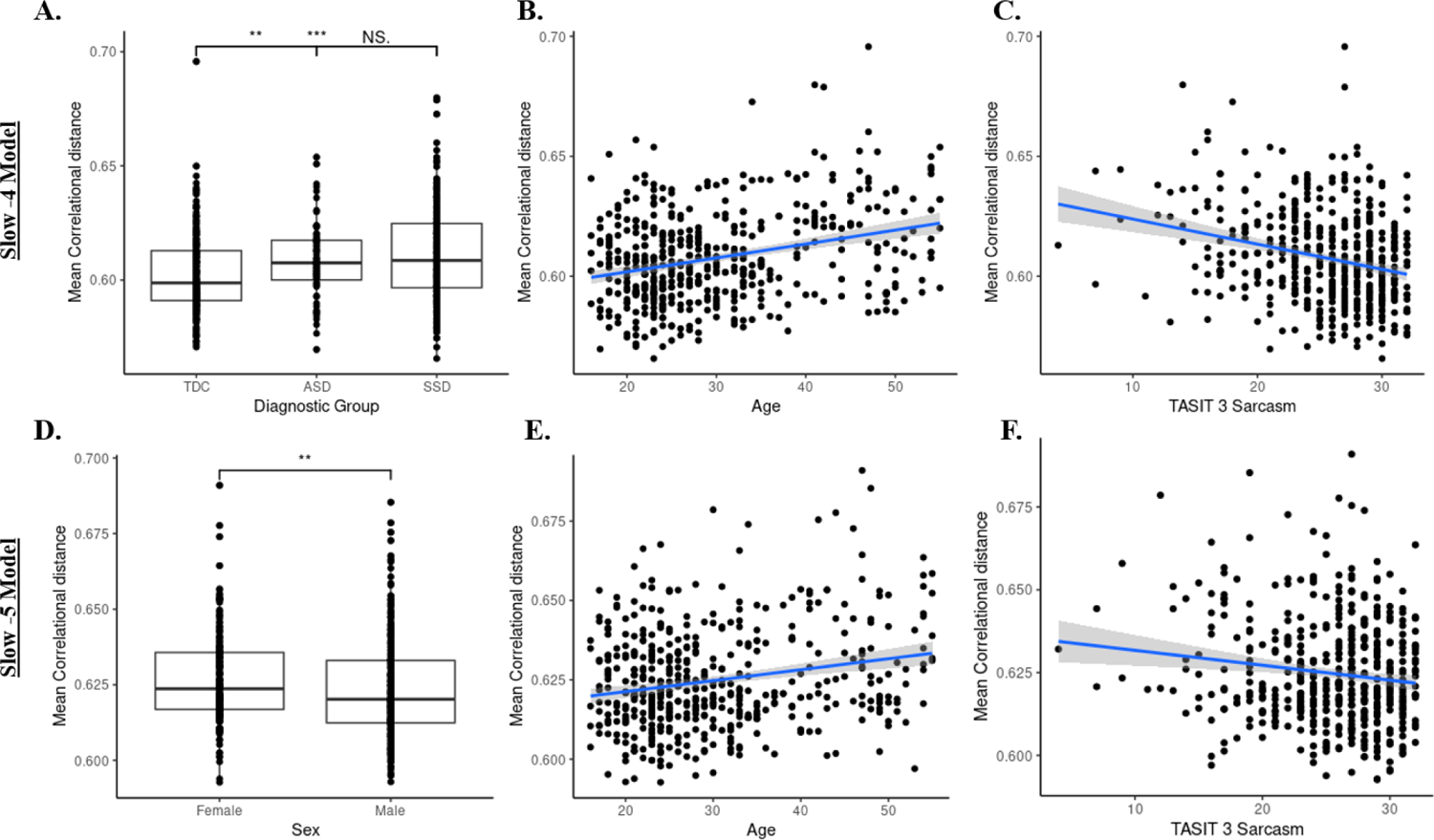
Mean correlational distance of fALFF slow-4 and slow-5 across participants. **A)** Boxplots demonstrate between-diagnostic group differences in mean correlational distance of fALFF slow-4 maps. The SSD group had a marginally higher mean correlational distance than the ASD and TDC. Dots represent individual data points. (‘NS’ p>0.1, ‘***’p<0.001, ‘**’ p<0.01) **B)** Higher mean correlational distance of slow-4 fALFF maps was associated with older age. Dots represent individual data points. Ribbon surrounding the regression line indicates a 95% confidence interval. **C)** Higher mean correlational distance of slow-4 fALFF maps was associated with worse TASIT-3-Sar scores. **D)** Boxplots demonstrate sex differences in mean correlational distance of slow-5 fALFF maps. Females had significantly higher mean correlational distances than males. (‘**’p<0.01) **E)** Higher mean correlational distance of slow-5 fALFF maps was associated with older age. **F)** Higher mean correlational distance of slow-5 fALFF maps was associated with worse TASIT-3-Sar scores.

#### 3.5.2 fALFF slow-5

After accounting for mean FD (F(1,481) = 11.10, p < 0.01) and site effects (F(2,481) = 0.32, p = 0.73), MCD of fALFF slow-5 maps did not differ across diagnostic groups, F(2,481) = 1.67, p = 0.19. However, there was a significant effect of sex on MCD, with females on average demonstrating greater variability in their spatial patterns of slow-5 fALFF compared to males, F(1,481) = 8.09, p < 0.01 (Figure 4D). Additionally, older age was related to increased MCD, F(1,481) = 14.69, p < 0.001 (Figure 4E), and lower TASIT-3-Sar score was associated with increased MCD, F(1,481) = 5.07, p = 0.025 (Figure 4F). Similar to findings regarding individual variability in slow-4 fALFF, the effect of social cognition was not influenced by the diagnostic group, F(2,481) = 0.51, p = 0.60.

## 4. Discussion

In this study, we extended the previous literature by: (1) performing a groupwise comparison of fALFF, a measure of spontaneous resting-state local brain activity, in ASD, SSD and TDC, (2) examining associations between fALFF and social cognition, and (3) investigating inter-individual variability within each diagnostic group. Our findings indicate lower slow-4 and slow-5 fALFF for both ASD and SSD compared with TDC primarily in the visual regions, specifically cuneus and occipital cortex. Furthermore, the SSD group showed significant differences in sensorimotor and subcortical regions relative to TDC, which were not observed in ASD. In addition, significant differences in fALFF between ASD and SSD were only present for limited regions in the cuneus. TASIT-3-Sar score, a measure of higher-order social cognition, but not ER40 scores, a measure of lower-order social cognition, was associated with slow-4 fALFF in the posterior regions, including the lateral occipitotemporal and temporoparietal regions, and with slow-5 fALFF in the brainstem. Consistent with recent work on task-related fMRI activity [23,24], a higher within-group inter-individual variability in fALFF was observed in the ASD and SSD groups compared to TDC. Lower variability was also related to higher TASIT-3-Sar scores, further suggesting that a more ‘typical’ pattern of brain activity was associated with cognitive outcomes [23,25].

Broadly speaking, patterns of fALFF differences relative to controls were similar in SSD and ASD. The findings in the occipital regions were consistent with a previous meta-analysis that found lower fALFF in the right occipital lobe in SSD [12], and another in children with ASD or early-onset schizophrenia compared to TDC in the visual regions [46]. This suggests some common deficits in early visual processing regions in SSD [47–49] and ASD [50–52], though ASD showed reduced fALLF relative to SSD in secondary visual cortex. Likewise, both SSD and ASD showed increased fALFF in prefrontal and anterior temporal regions, though these results were less prominent in the age- and sex-matched analysis, suggesting that these effects may be somewhat influenced by age and/or sex. Within SSD, diagnostic specific differences from controls were noted in motor cortex, consistent with [12], and subcortical regions (caudate, striatum, hippocampal, and amygdala), but these regions did not significantly differ between SSD and ASD. Subcortical circuits involving the striatum have been identified as impaired in SSD [53], and corticostriatal connectivity has been linked to treatment outcomes in SSD [54,55]. The hippocampus and amygdala have been identified as the subcortical structures with the greatest volume reduction in SSD [56], whereas subcortical volume differences appear to be less pronounced in ASD [57]. The analysis of fALLF across SSD, ASD, and TDC suggest that while there may be some regional differences in magnitude of fALFF between SSD and ASD relative to controls, a broadly similar pattern of deficits is present across both disorders, with decreased fALFF in sensory-motor and increased fALFF in prefrontal cortices.

We found that across the whole sample, higher-level social cognition as measured by TASIT-3-Sar scores was positively correlated with higher fALFF in the temporoparietal and lateral occipitotemporal junction, which is in line with the overall body of literature on social cognition, primarily examining higher-level social cognition in TDC [58–61] individuals with SSD [58,59] and ASD [62]. Furthermore, while we did not find a main effect of low-level social cognition (i.e. ER40) on fALFF in the overall sample, our findings pertaining to the interaction of fALFF and ER40 scores in ASD compared with SSD and TDC suggest the involvement of early visual regions, such as cuneus, in emotional recognition; this interaction effect indicated that as ER40 scores increased, lower fALFF in early visual regions were observed in ASD but higher fALFF were observed in SSD and TDC. This result is consistent with previous literature that showed that substantial differences exist in how social information is processed in the early visual cortex for ASD compared with other healthy and clinical populations [63].

The increased variability in fALFF in SSD and ASD is a key finding consistent with a growing body of literature noting greater heterogeneity of functional brain activation/connectivity patterns in psychiatric samples [24,36,64–69]. Mean correlational distance quantifies how “typical”, or close to the average, one’s pattern of brain activity is. One benefit of this approach is that it does not presuppose that all participants in a given diagnostic group share a common spatial pattern of deficits; this is particularly important given the high heterogeneity in prior work on fALFF in SSD [12] and ASD [70].

Interestingly, these differences were only present in slow-4, highlighting the importance of considering the different frequency bands within fALFF [8]. In addition, increased variability was also associated with lower TASIT-3-Sar scores, highlighting that more typical fALFF patterns may be beneficial to social cognitive performance. Diagnostic differences in inter-individual variability highlight the importance of moving beyond comparing the average values in clinical populations and TDC in spatially specific regions, given the heterogeneity observed in clinical groups [66]. Instead, variability may represent a novel biomarker that could also be used as a metric of treatment response, without presupposing that every patient shares common neurobiological deficits or common treatment responses.

Despite the large sample size, this study was not without its limitations. A short resting-state scan of 7 minutes was used; a longer scan duration could produce more reliable results [71,72] with increased sensitivity to detect group differences. The mean age of ASD participants was different from that of SSD and TDC groups in the full sample, and while protocols were harmonized, the ASD group was collected at a single site as part of a parallel study. This may have introduced sampling biases in the recruitment of ASD versus SSD. The SPIN-ASD dataset was also partially collected during the COVID-19 pandemic when social interactions were heavily restricted, which may have affected who was willing to participate in the study as well as overall social activities, captured using social functioning scales, such as the BSFS. The ASD sample also consisted of a smaller sample of individuals with ASD, and may not have included the full range of social cognitive impairments observed in SSD.

This was the largest study to date incorporating both SSD and ASD while examining fALFF and social cognition. ASD and SSD showed broadly similar patterns, with common deficits of fALFF in the occipital cortex. The differences in individual variability as measured by MCD highlight a growing perspective that rather than focusing on common diagnostic group differences, individuals with psychiatric disorders can demonstrate highly heterogeneous deviations in brain structure or function [66]. A ‘normative’ modeling approach, moving away from group-linear approaches and considering variability and deviation from the norm across multiple systems provides a new avenue for discovery in psychiatric neuroscience.

## Acknowledgments

The authors acknowledge the important contributions of the SPINS and SPIN-ASD teams for the collection of both data sets, as well as the staff and participants who made these studies possible.

## Author Contribution

- SB led the study by data analysis, study idea conceptualization and drafting and preparing the manuscript.
- JCY assisted in data analysis and preparing the manuscript.
- JG and LDO assisted in data analysis, study idea conceptualization and preparing the manuscript.
- VT and AGR assisted in data analysis.
- EWD assisted by providing the study data, conceptualization and data analysis.
- GF, MCL, RWB, AKM, ANV and SHA contributed by leading data acquisition, providing data and assisting in study conceptualization.
- CH conceptualized the study ideas and prepared the manuscript.
- SHA and CH provided major supervision throughout the project.
- All authors reviewed and approved the final version of the manuscript.

## References

1. Zeidan J, Fombonne E, Scorah J, Ibrahim A, Durkin MS, Saxena S, et al. Global prevalence of autism: A systematic review update. Autism Res. 2022;15:778–790.

2. Kahn RS, Sommer IE, Murray RM, Meyer-Lindenberg A, Weinberger DR, Cannon TD, et al. Schizophrenia (Primer). Nature Reviews: Disease Primers. 2015;1.

3. Yu-Feng Z, Yong H, Chao-Zhe Z, Qing-Jiu C, Man-Qiu S, Meng L, et al. Altered baseline brain activity in children with ADHD revealed by resting-state functional MRI. Brain and Development. 2007;29:83–91.

4. Zou Q-H, Zhu C-Z, Yang Y, Zuo X-N, Long X-Y, Cao Q-J, et al. An improved approach to detection of amplitude of low-frequency fluctuation (ALFF) for resting-state fMRI: fractional ALFF. J Neurosci Methods. 2008;172:137–141.

5. Hoptman MJ, Zuo X-N, Butler PD, Javitt DC, D’Angelo D, Mauro CJ, et al. Amplitude of low-frequency oscillations in schizophrenia: a resting state fMRI study. Schizophr Res. 2010;117:13– 20.

6. Xu Y, Zhuo C, Qin W, Zhu J, Yu C. Altered Spontaneous Brain Activity in Schizophrenia: A Meta-Analysis and a Large-Sample Study. Biomed Res Int. 2015;2015:204628.

7. Zhou C, Tang X, You W, Wang X, Zhang X, Zhang X, et al. Altered Patterns of the Fractional Amplitude of Low-Frequency Fluctuation and Functional Connectivity Between Deficit and Non-Deficit Schizophrenia. Front Psychiatry. 2019;10:680.

8. Yu R, Chien Y-L, Wang H-LS, Liu C-M, Liu C-C, Hwang T-J, et al. Frequency-specific alternations in the amplitude of low-frequency fluctuations in schizophrenia. Hum Brain Mapp. 2014;35:627–637.

9. Alonso-Solís A, Vives-Gilabert Y, Portella MJ, Rabella M, Grasa EM, Roldán A, et al. Altered amplitude of low frequency fluctuations in schizophrenia patients with persistent auditory verbal hallucinations. Schizophr Res. 2017;189:97–103.

10. Meda SA, Wang Z, Ivleva EI, Poudyal G, Keshavan MS, Tamminga CA, et al. Frequency-Specific Neural Signatures of Spontaneous Low-Frequency Resting State Fluctuations in Psychosis: Evidence From Bipolar-Schizophrenia Network on Intermediate Phenotypes (B-SNIP) Consortium. Schizophr Bull. 2015;41:1336–1348.

11. Liu S, Guo Z, Cao H, Li H, Hu X, Cheng L, et al. Altered asymmetries of resting-state MRI in the left thalamus of first-episode schizophrenia. Chronic Dis Transl Med. 2022;8:207–217.

12. Gong J, Wang J, Luo X, Chen G, Huang H, Huang R, et al. Abnormalities of intrinsic regional brain activity in first-episode and chronic schizophrenia: a meta-analysis of resting-state functional MRI. J Psychiatry Neurosci. 2020;45:55–68.

13. Li G, Rossbach K, Jiang W, Du Y. Resting-state brain activity in Chinese boys with low functioning autism spectrum disorder. Ann Gen Psychiatry. 2018;17:47.

14. Karavallil Achuthan S, Coburn KL, Beckerson ME, Kana RK. Amplitude of low frequency fluctuations during resting state fMRI in autistic children. Autism Res. 2023;16:84–98.

15. Sasson NJ, Pinkham AE, Carpenter KLH, Belger A. The benefit of directly comparing autism and schizophrenia for revealing mechanisms of social cognitive impairment. J Neurodev Disord. 2011;3:87–100.

16. Beaudoin C, Beauchamp MH. Social cognition. Handb Clin Neurol. 2020;173:255–264.

17. Oliver, Haltigan, Gold, Foussias. Lower-and higher-level social cognitive factors across individuals with schizophrenia spectrum disorders and healthy controls: relationship with neurocognition and …. Schizophr Res. 2019. 2019.

18. Green MF, Horan WP, Lee J. Social cognition in schizophrenia. Nat Rev Neurosci. 2015;16:620– 631.

19. Oliver LD, Moxon-Emre I, Lai M-C, Grennan L, Voineskos AN, Ameis SH. Social Cognitive Performance in Schizophrenia Spectrum Disorders Compared With Autism Spectrum Disorder: A Systematic Review, Meta-analysis, and Meta-regression. JAMA Psychiatry. 2021;78:281–292.

20. Sui J, Pearlson GD, Du Y, Yu Q, Jones TR, Chen J, et al. In search of multimodal neuroimaging biomarkers of cognitive deficits in schizophrenia. Biol Psychiatry. 2015;78:794–804.

21. Guo P, Hu S, Jiang X, Zheng H, Mo D, Cao X, et al. Associations of Neurocognition and Social Cognition With Brain Structure and Function in Early-Onset Schizophrenia. Front Psychiatry. 2022;13:798105.

22. Jung M, Tu Y, Lang CA, Ortiz A, Park J, Jorgenson K, et al. Decreased structural connectivity and resting-state brain activity in the lateral occipital cortex is associated with social communication deficits in boys with autism spectrum disorder. Neuroimage. 2019;190:205–212.

23. Hawco C, Yoganathan L, Voineskos AN, Lyon R, Tan T, Daskalakis ZJ, et al. Greater Individual Variability in Functional Brain Activity during Working Memory Performance in young people with Autism and Executive Function Impairment. Neuroimage Clin. 2020;27:102260.

24. Gallucci J, Tan T, Schifani C, Dickie EW, Voineskos AN, Hawco C. Greater individual variability in functional brain activity during working memory performance in Schizophrenia Spectrum Disorders (SSD). Schizophr Res. 2022;248:21–31.

25. Gallucci J, Pomarol-Clotet E, Voineskos AN, Guerrero-Pedraza A, Alonso-Lana S, Vieta E, et al. Longer illness duration is associated with greater individual variability in functional brain activity in Schizophrenia, but not bipolar disorder. Neuroimage Clin. 2022;36:103269.

26. Viviano JD, Buchanan RW, Calarco N, Gold JM, Foussias G, Bhagwat N, et al. Resting-State Connectivity Biomarkers of Cognitive Performance and Social Function in Individuals With Schizophrenia Spectrum Disorder and Healthy Control Subjects. Biol Psychiatry. 2018;84:665– 674.

27. Oliver LD, Hawco C, Homan P, Lee J, Green MF, Gold JM, et al. Social cognitive networks and social cognitive performance across individuals with schizophrenia spectrum disorders and healthy controls. Biological Psychiatry: Cognitive Neuroscience and Neuroimaging. 2020. 2020.

28. Lord C, Rutter M, DiLavore PC, Risi S. Autism Diagnostic Observation Schedule: Ados-2. Western Psychological Services; 2006.

29. Wechsler D. Wechsler Test of Adult Reading: WTAR. Psychological Corporation; 2001.

30. Overall JE, Gorham DR. The Brief Psychiatric Rating Scale. Psychol Rep. 1962;10:799–812.

31. Andreasen NC. Negative symptoms in schizophrenia. Definition and reliability. Arch Gen Psychiatry. 1982;39:784–788.

32. Buchanan RW, Javitt DC, Marder SR, Schooler NR, Gold JM, McMahon RP, et al. The Cognitive and Negative Symptoms in Schizophrenia Trial (CONSIST): the efficacy of glutamatergic agents for negative symptoms and cognitive impairments. Am J Psychiatry. 2007;164:1593–1602.

33. Kohler CG, Bilker W, Hagendoorn M, Gur RE, Gur RC. Emotion recognition deficit in schizophrenia: association with symptomatology and cognition. Biol Psychiatry. 2000;48:127–136.

34. McDonald S, Flanagan S, Martin I, Saunders C. The ecological validity of TASIT: A test of social perception. Neuropsychol Rehabil. 2004;14:285–302.

35. Mancuso F, Horan WP, Kern RS, Green MF. Social cognition in psychosis: multidimensional structure, clinical correlates, and relationship with functional outcome. Schizophr Res. 2011;125:143–151.

36. Hawco C, Buchanan RW, Calarco N, Mulsant BH, Viviano JD, Dickie EW, et al. Separable and Replicable Neural Strategies During Social Brain Function in People With and Without Severe Mental Illness. Am J Psychiatry. 2019:appiajp201817091020.

37. Hawco C, Dickie EW, Herman G, Turner JA, Argyelan M, Malhotra AK, et al. A longitudinal multi-scanner multimodal human neuroimaging dataset. Sci Data. 2022;9:332.

38. Dickie EW, Anticevic A, Smith DE, Coalson TS, Manogaran M, Calarco N, et al. ciftify: A framework for surface-based analysis of legacy MR acquisitions. bioRxiv. 2018:484428.

39. Xue S-W, Li D, Weng X-C, Northoff G, Li D-W. Different neural manifestations of two slow frequency bands in resting functional magnetic resonance imaging: a systemic survey at regional, interregional, and network levels. Brain Connect. 2014;4:242–255.

40. Zuo X-N, Di Martino A, Kelly C, Shehzad ZE, Gee DG, Klein DF, et al. The oscillating brain: complex and reliable. Neuroimage. 2010;49:1432–1445.

41. Radua J, Vieta E, Shinohara R, Kochunov P, Quidé Y, Green MJ, et al. Increased power by harmonizing structural MRI site differences with the ComBat batch adjustment method in ENIGMA. Neuroimage. 2020;218:116956.

42. Yu M, Linn KA, Cook PA, Phillips ML, McInnis M, Fava M, et al. Statistical harmonization corrects site effects in functional connectivity measurements from multi-site fMRI data. Hum Brain Mapp. 2018;39:4213–4227.

43. Smith SM, Nichols TE. Threshold-free cluster enhancement: addressing problems of smoothing, threshold dependence and localisation in cluster inference. Neuroimage. 2009;44:83–98.

44. Power JD, Mitra A, Laumann TO, Snyder AZ, Schlaggar BL, Petersen SE. Methods to detect, characterize, and remove motion artifact in resting state fMRI. NeuroImage. 2014;84:320–341.

45. Leucht S, Samara M, Heres S, Davis JM. Dose Equivalents for Antipsychotic Drugs: The DDD Method. Schizophr Bull. 2016;42 Suppl 1:S90–S94.

46. Jin X, Zhang K, Lu B, Li X, Yan C-G, Du Y, et al. Shared atypical spontaneous brain activity pattern in early onset schizophrenia and autism spectrum disorders: evidence from cortical surface-based analysis. Eur Child Adolesc Psychiatry. 2023. 26 December 2023. 10.1007/s00787-023-02333-2.

47. Martínez A, Gaspar PA, Bermudez DH, Aburto-Ponce MB, Javitt DC. Bottom-up and top-down contributions to impaired motion processing in schizophrenia. medRxiv. 2023. 8 July 2023. 10.1101/2023.07.07.23292259.

48. Martino M, Magioncalda P. A working model of neural activity and phenomenal experience in psychosis. Mol Psychiatry. 2024. 6 June 2024. 10.1038/s41380-024-02607-4.

49. Schoonover KE, Dienel SJ, Holly Bazmi H, Enwright JF 3rd, Lewis DA. Altered excitatory and inhibitory ionotropic receptor subunit expression in the cortical visuospatial working memory network in schizophrenia. Neuropsychopharmacology. 2024;49:1183–1192.

50. Jassim N, Baron-Cohen S, Suckling J. Meta-analytic evidence of differential prefrontal and early sensory cortex activity during non-social sensory perception in autism. Neurosci Biobehav Rev. 2021;127:146–157.

51. Vandewouw MM, Choi E, Hammill C, Arnold P, Schachar R, Lerch JP, et al. Emotional face processing across neurodevelopmental disorders: a dynamic faces study in children with autism spectrum disorder, attention deficit hyperactivity disorder and obsessive-compulsive disorder. Transl Psychiatry. 2020;10:375.

52. Degré-Pelletier J, Danis É, Thérien VD, Bernhardt B, Barbeau EB, Soulières I. Differential neural correlates underlying visuospatial versus semantic reasoning in autistic children. Cereb Cortex. 2024;34:19–29.

53. Ji JL, Diehl C, Schleifer C, Tamminga CA, Keshavan MS, Sweeney JA, et al. Schizophrenia Exhibits Bi-directional Brain-Wide Alterations in Cortico-Striato-Cerebellar Circuits. Cereb Cortex. 2019;29:4463–4487.

54. Sarpal DK, Robinson DG, Lencz T, Argyelan M, Ikuta T, Karlsgodt K, et al. Antipsychotic treatment and functional connectivity of the striatum in first-episode schizophrenia. JAMA Psychiatry. 2015;72:5–13.

55. Sarpal DK, Argyelan M, Robinson DG, Szeszko PR, Karlsgodt KH, John M, et al. Baseline Striatal Functional Connectivity as a Predictor of Response to Antipsychotic Drug Treatment. Am J Psychiatry. 2016;173:69–77.

56. van Erp TGM, Hibar DP, Rasmussen JM, Glahn DC, Pearlson GD, Andreassen OA, et al. Subcortical brain volume abnormalities in 2028 individuals with schizophrenia and 2540 healthy controls via the ENIGMA consortium. Mol Psychiatry. 2016;21:547–553.

57. van Rooij D, Anagnostou E, Arango C, Auzias G, Behrmann M, Busatto GF, et al. Cortical and Subcortical Brain Morphometry Differences Between Patients With Autism Spectrum Disorder and Healthy Individuals Across the Lifespan: Results From the ENIGMA ASD Working Group. Am J Psychiatry. 2018;175:359–369.

58. Schurz M, Radua J, Aichhorn M, Richlan F, Perner J. Fractionating theory of mind: a meta-analysis of functional brain imaging studies. Neurosci Biobehav Rev. 2014;42:9–34.

59. Kronbichler L, Tschernegg M, Martin AI, Schurz M, Kronbichler M. Abnormal Brain Activation During Theory of Mind Tasks in Schizophrenia: A Meta-Analysis. Schizophr Bull. 2017;43:1240– 1250.

60. Tso IF, Rutherford S, Fang Y, Angstadt M, Taylor SF. The ‘social brain’ is highly sensitive to the mere presence of social information: An automated meta-analysis and an independent study. PLoS One. 2018;13:e0196503.

61. Van Overwalle F. Social cognition and the brain: a meta-analysis. Hum Brain Mapp. 2009;30:829– 858.

62. Hao Z, Shi Y, Huang L, Sun J, Li M, Gao Y, et al. The Atypical Effective Connectivity of Right Temporoparietal Junction in Autism Spectrum Disorder: A Multi-Site Study. Front Neurosci. 2022;16:927556.

63. Mottron L, Dawson M, Soulières I, Hubert B, Burack J. Enhanced perceptual functioning in autism: an update, and eight principles of autistic perception. J Autism Dev Disord. 2006;36:27–43.

64. Dickie EW, Ameis SH, Shahab S, Calarco N, Smith DE, Miranda D, et al. Personalized Intrinsic Network Topography Mapping and Functional Connectivity Deficits in Autism Spectrum Disorder. Biol Psychiatry. 2018;84:278–286.

65. Dickie EW, Shahab S, Hawco C, Miranda D, Herman G, Argyelan M, et al. Robust hierarchically organized whole-brain patterns of dysconnectivity in schizophrenia spectrum disorders observed after Personalized Intrinsic Network Topography. bioRxiv. 2022:2022.12.13.520333.

66. Segal A, Parkes L, Aquino K, Kia SM, Wolfers T, Franke B, et al. Regional, circuit and network heterogeneity of brain abnormalities in psychiatric disorders. Nat Neurosci. 2023;26:1613–1629.

67. Chen H, Nomi JS, Uddin LQ, Duan X, Chen H. Intrinsic functional connectivity variance and state-specific under-connectivity in autism. Hum Brain Mapp. 2017;38:5740–5755.

68. Hahamy A, Behrmann M, Malach R. The idiosyncratic brain: distortion of spontaneous connectivity patterns in autism spectrum disorder. Nat Neurosci. 2015;18:302–309.

69. Gopal S, Miller RL, Michael A, Adali T, Cetin M, Rachakonda S, et al. Spatial Variance in Resting fMRI Networks of Schizophrenia Patients: An Independent Vector Analysis. Schizophr Bull. 2016;42:152–160.

70. Mei T, Ma Z-H, Guo Y-Q, Lu B, Cao Q-J, Chen X, et al. Frequency-specific age-related changes in the amplitude of spontaneous fluctuations in autism. Transl Pediatr. 2022;11:349–358.

71. Gordon EM, Laumann TO, Gilmore AW, Newbold DJ, Greene DJ, Berg JJ, et al. Precision Functional Mapping of Individual Human Brains. Neuron. 2017;95:791–807.e7.

72. Noble S, Scheinost D, Constable RT. A decade of test-retest reliability of functional connectivity: A systematic review and meta-analysis. Neuroimage. 2019;203:116157.

