## Supplemental Figures 1 and 2 for "Transdiagnostic Neurobiology of Social Cognition and Individual Variability as Measured by Fractional Amplitude of Low-Frequency Fluctuation in Schizophrenia and Autism Spectrum Disorders"

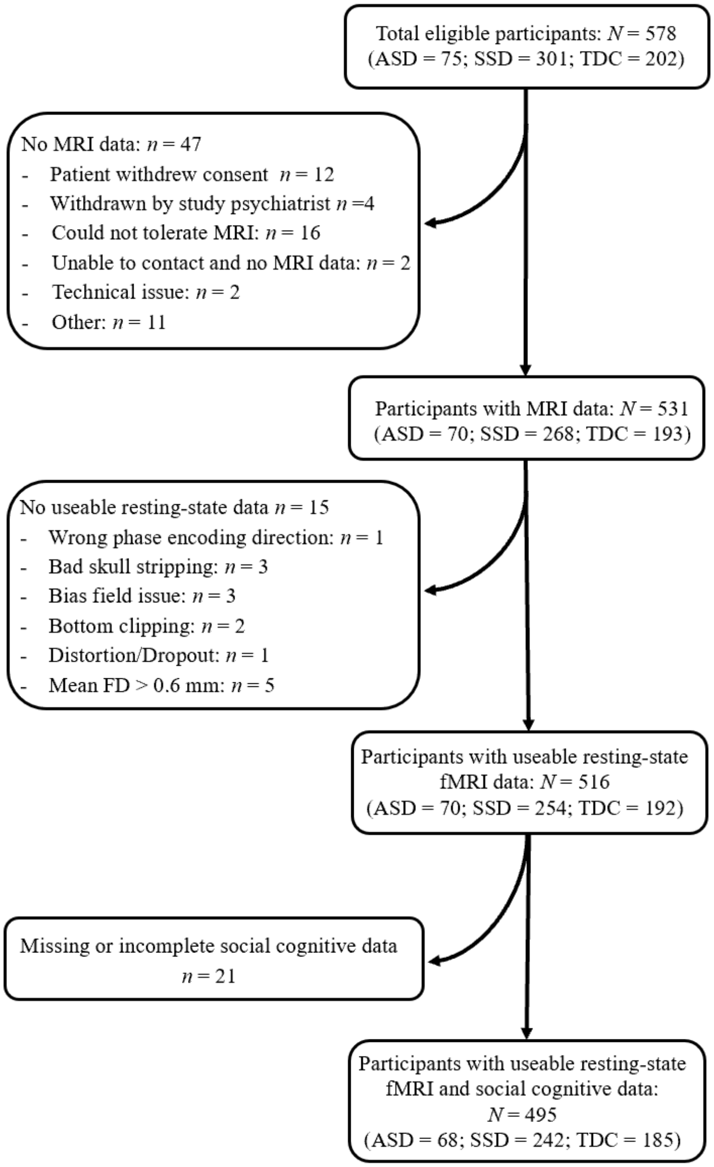


**Supplemental Figure 1:** Consort diagram indicating the total number of participants eligible for the study and those with usable resting-state and social cognitive performance data.

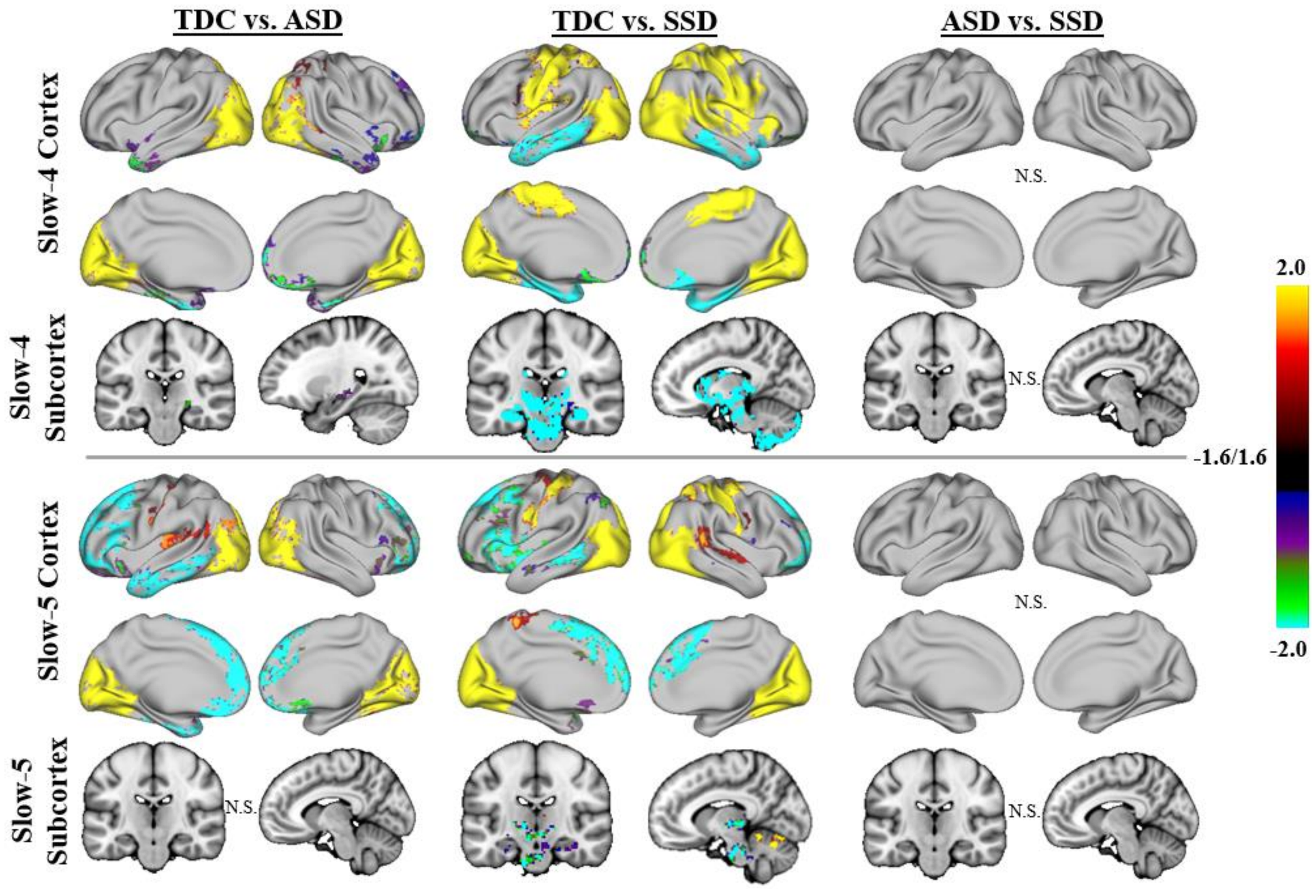


**Supplemental Figure 2:** Group results for linear models examining fALFF among ASD, SSD and TDC without TASIT-3-Sar and ER40 scores included as covariates in the model. Similar to Figure 1, results of whole-brain, vertex-wise analysis using TFCE are presented, in brain-wide FWE corrected log-10(p), thresholded at 1.6, equal to p = 0.025 (to correct for left and right hemisphere). TDC showed significantly higher slow-4 and slow-5 fALFF than both ASD and SSD in the visual regions (i.e. cuneus and occipital cortex; yellow-red regions). In addition, TDC also had higher fALFF than SSD in the sensory-motor cortex (yellow-red regions). Increased slow-5 fALFF was observed in both ASD and SSD compared with TDC in the medial frontal regions (green-blue). SSD also showed greater fALFF than TDC in the subcortical regions (green-blue). No significant differences in slow-4 or slow-5 fALFF were observed between ASD and SSD.
